# Hepatitis E Virus infection and waste pickers: a case-control seroprevalence study

**DOI:** 10.1101/679076

**Authors:** Cosme Alvarado-Esquivel, Verónica Dayali Gutierrez-Martinez, Eda Guadalupe Ramírez-Valles, Antonio Sifuentes-Alvarez

## Abstract

Whether waste pickers are a risk group for hepatitis E virus (HEV) infection is largely unknown. Therefore, this study aimed to determine the association between HEV exposure and: 1) the occupation of waste picker; and 2) the work characteristics of waste pickers. An age-and gender-matched case-control seroprevalence study of 86 waste pickers (mean age: 35.45 ± 17.15 years) and 86 control subjects of the general population was performed. We determined anti-HEV IgG antibodies in sera of cases and controls using a commercially available enzyme-linked immunoassay. The McNemar’s test was used to assess the association between HEV seropositivity and the occupation of waste picker. The association between HEV seropositivity and work characteristics of waste pickers was assessed by bivariate and regression analyses. Anti-HEV IgG antibodies were detected in 14 (16.3%) of the 86 waste pickers and in 8 (9.3%) of the 86 control subjects (McNemar’s pair test: OR = 13.0; 95% CI: 0.73-230.77; *P*=0.02). Ill waste pickers had a higher HEV seroprevalence than those who were apparently healthy (6/15: 40% vs 8/71: 11.3%, respectively: *P*=0.01). Waste pickers with reflexes impairment had a higher HEV seroprevalence than those without this impairment (5/10: 50% and 9/76: 11.8%, respectively; *P*=0.009). Logistic regression analysis of sociodemographic, work, and behavioral characteristics of waste pickers showed that HEV seropositivity was associated with increasing age (OR = 6.52; 95% CI: 1.95-21.78; *P*=0.002) and raising pigs (OR = 12.01; 95% CI: 1.48-97.26; *P*=0.02). This is the first age- and gender-matched case-control study on the association between HEV infection and the occupation of waste picker. Waste pickers represent a risk group for HEV infection. Factors associated with HEV seropositivity found in this study may help in the design of optimal planning to avoid HEV infection.

## Introduction

Estimates indicate that there are about 20 million cases of acute hepatitis E and 56000 deaths associated with this disease in the world every year [1]. Hepatitis E results from infection with hepatitis E virus that is a single-stranded, positive sense RNA virus [2]. Predominant route of infection with HEV varies depending on the development of the country. In developing countries, HEV transmission occurs via contaminated water and food by the fecal oral route; whereas in developed countries, this infection occurs by zoonotic transmission [3]. Other transmission routes of HEV include blood transfusion [4], hemodialysis [5], and solid organ transplantation [6]. Chronic infections with HEV have been reported in immunocompromised patients, and a high seroprevalence of HEV was found in patients with autoimmune hepatitis [7]. Fulminant hepatitis E has been reported in pregnant women, patients with underlying liver disease [8], and children [9]. Hepatitis E may present as sporadic cases or outbreaks [10]. Consumption of meat from HEV-infected animals including, for instance, pigs and boars may lead to infection in humans [11].

Transmission of HEV occurs in areas where poor sanitation and weak public health infrastructure exist [12]. Waste pickers is a group of people in close contact with garbage, dirty food and liquids in their occupational setting. Sometimes, they even eat food from the garbage. We are aware of only one study on the seroepidemiology of HEV infection in waste pickers. In a cross-sectional study in Brazil, researchers found a 5.1% seroprevalence of HEV infection in waste pickers [13]. Whether waste picking occupation is a risk for HEV infection is largely unknown. Therefore, we aimed to determine the association between anti-HEV IgG antibodies seropositivity and: 1) the occupation of waste picker; and 2) the work characteristics of waste pickers.

## Materials and Methods

### Study design and populations studied

Through a case-control study design 86 waste pickers (cases) and 86 age- and gender matched control subjects without waste picker occupation (controls) were studied. Inclusion criteria for enrollment of waste pickers were: 1) subjects with an occupation of waste picker in Durango City, Mexico; 2) 14 years and older; and 3) who accepted to participate in the study. Gender and socioeconomic level were not restrictive criteria for enrollment of cases. Fifty-four (62.8%) of the waste pickers were females and 32 (37.2%) were males. Cases were 14 to 76 (mean: 35.45 ± 17.15) years old. On the other hand, inclusion criteria for enrollment of control subjects included: 1) individuals without waste picker occupation of the general population from Durango City; 2) 14 years and older; and 3) who accepted to participate in the study. The control group included 54 (62.8%) females and 32 (37.2%) males. Controls were 16 to 78 (mean: 36.05 ± 17.02) years old. Gender and age were comparable in cases and controls (*P*=1.00 and *P*=0.81, respectively).

### Characteristics of waste pickers

We obtained the socio-demographic, clinical, work, housing and behavioral characteristics of waste pickers with the aid of a questionnaire. Sociodemographic items included gender, age, birthplace, residence, education, and socioeconomic level. Clinical items included the self-reported health status (ill or healthy), history of blood transfusion, and impairments in memory, reflexes, hearing and vision. Work characteristics included seniority, history of injuries with sharp material, wearing hand gloves and face masks, eating or drinking alcohol while working, and eating from the garbage. In addition, a habitual washing hands before eating was recorded. With respect to behavioral data, the characteristics drinking untreated water or unpasteurized milk, consumption of unwashed raw vegetables or fruits, consumption of raw or undercooked meat, type of meat consumed (pork, boar, rabbit, beef, goat, lamb, venison, squirrel, horse, rat, chicken, turkey, and fish), consumption of sausages, ham, salami or chorizo, contact with animals, traveling, and contact with soil (gardening or agriculture). As to housing characteristics, we used the Bronfman’s criteria [14]: number of rooms in the house, number of persons in the house, availability of drinkable water, form of elimination of excreta, and type of flooring (ceramic, wood, concrete, soil).

### Detection of anti-HEV IgG antibodies

We obtained a blood sample from each subject. Serum was obtained from the blood sample by centrifugation. Serum samples were frozen at −20°C until analyzed. Detection of anti-HEV IgG antibodies in serum samples was performed by using a commercially available enzyme immunoassay kit: “AccuDiag™ HEV IgG ELISA” (Diagnostic Automation Inc., Woodland Hills, CA. USA). All assays were performed following the manufacturer’s instructions.

### Statistical Analysis

The statistical analysis was performed with the software Microsoft Excel, Epi Info version 7, and SPSS version 20. For the sample size calculation, we used the following parameters: a 95% two-sided confidence level, a 1:1 ratio of cases and controls, a power of 80%, a 36.6% outcome in unexposed group [15], and an odds ratio (OR) of 2.4. The result of the sample size calculation was 84 cases and 84 controls. Age in cases and controls was compared using the student’s *t* test. HEV IgG seropositivity rates in cases and controls were compared using the McNemar’s paired test. We used the Pearson’s chi-squared test or the two-tailed Fisher’s exact test (for values <5) to assess the association between the HEV IgG seropositivity rate and the characteristics of waste pickers. Then, socio-demographic, work, housing, and behavioral characteristics of waste pickers with a *P* value < 0.05 obtained in the bivariate analysis were selected for further analysis using regression analysis with the Enter method. We calculated the odds ratio (OR) and 95% confidence interval (CI), and a *P* value < 0.05 was considered as statistically significant.

### Ethical aspects

The Ethical Committee of the Faculty of Medicine and Nutrition of the Juárez University of Durango State in Durango City, Mexico approved this study. Participants were informed about the aims, and procedures of the study. Participation in the study was voluntary, and a written informed consent was obtained from all subjects and the next of kin of minor participants.

## Results

Anti-HEV IgG antibodies were detected in 14 (16.3%) of the 86 waste pickers and in 8 (9.3%) of the 86 control subjects. The difference between the HEV seroprevalences in cases and controls was statistically significant according to the McNemar’s pair test (OR = 13.0; 95% CI: 0.73-230.77; *P*=0.02) (Table 1). As to the correlation of socio-demographic characteristics of waste pickers and HEV seropositivity, bivariate analysis showed that HEV seroprevalence increased with age (*P*<0.001). Other sociodemographic characteristics of waste pickers including gender, residence, socio-economic status or educational level did not show an association with HEV seroprevalence. A correlation between HEV seropositivity and sociodemographic characteristics of waste pickers is shown in Table 2. Concerning clinical characteristics, waste pickers who referred themselves as ill had a higher HEV seroprevalence than those who were apparently healthy (6/15: 40% vs 8/71: 11.3%, respectively: *P*=0.01). HEV seroprevalence was comparable (*P*=0.63) in waste pickers with a history of blood transfusion (2/9: 22.2%) than in those without blood transfusion (12/77: 15.6%). Waste pickers with reflexes impairment had a higher HEV seroprevalence than those without this impairment (5/10: 50% and 9/76: 11.8%, respectively; *P*=0.009). Impairments in memory, hearing and vision did not show an association with HEV seropositivity. Of the work characteristics, HEV seroprevalence increased with seniority (Table 3), whereas other work characteristics including history of injuries with sharp material, wearing hand gloves and face masks, eating or drinking alcohol while working, washing hands before eating, and eating from the garbage did not show an association with HEV seropositivity. As to behavioral characteristics of waste pickers, bivariate analysis showed that raising pigs, consumption of venison, and squirrel meat were associated (*P*<0.05) with HEV seropositivity (Table 4). Other behavioral characteristics of waste pickers including consumption of untreated water, unpasteurized milk, unwashed raw vegetables or fruits, raw or undercooked meat, consumption of meat other than venison or squirrel meat, consumption of sausages, ham, salami or chorizo, contact with animals other than pigs, traveling and contact with soil did not show an association with HEV seropositivity. None of the housing characteristics of waste pickers including number of rooms in the house, number of persons in the house, availability of drinkable water, form of elimination of excreta, and type of flooring were associated with HEV seropositivity. Further analysis by logistic regression of sociodemographic, work, and behavioral characteristics of waste pickers with a *P*<0.05 obtained in bivariate analysis showed that HEV seropositivity was associated with increasing age (OR = 6.52; 95% CI: 1.95-21.78; *P*=0.002) and raising pigs (OR = 12.01; 95% CI: 1.48-97.26; *P*=0.02) (Table 5).

**Table 1:** Distribution of HEV infection among case control pairs.

| **Case** | **Control** |  | **Total** | **OR [95% CI]** | **P-value** |
| --- | --- | --- | --- | --- | --- |
|  | Exposed<br>(HEV<br>infection) | Unexposed<br>(No HEV<br>infection) |  |  |  |
| Exposed<br>(HEV<br>infection) | 8 | 6 | 14 |  |  |
| Unexposed<br>(No HEV<br>infection) | 0 | 72 | 72 |  |  |
| **Total** | 8 | 78 | 86 | 13.0 [0.73-230.77] | 0.02 |

0.5 has been added to each cell for calculation.

**Table 2.**
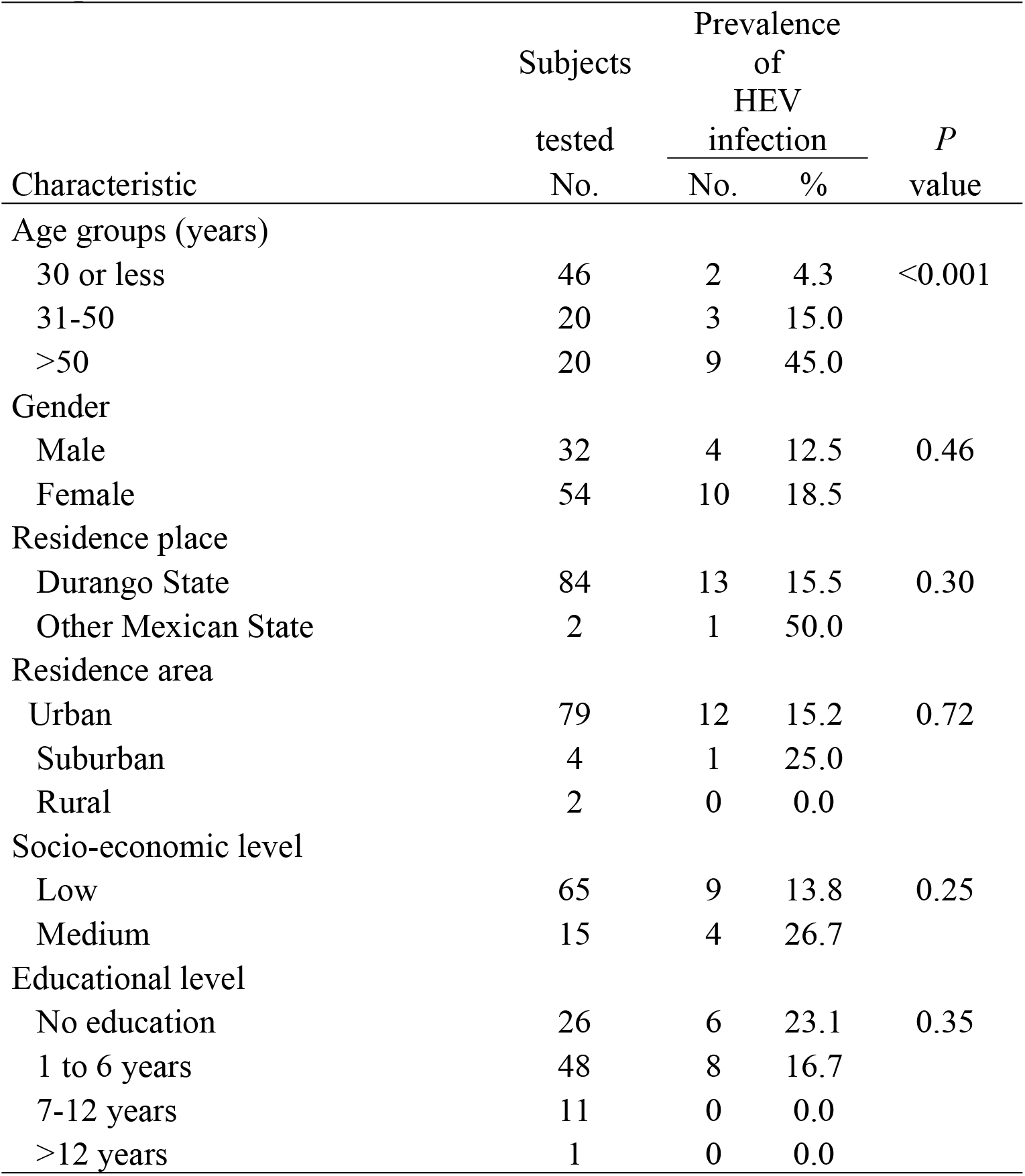
Socio-demographic characteristics of waste pickers and prevalence of HEV infection.

| Characteristic | Subjects<br>tested<br>No. | Prevalence<br>of<br>HEV<br>infection |  | *P*<br>value |
| --- | --- | --- | --- | --- |
|  |  | No. | % |  |
| Age groups (years) |  |  |  |  |
| 30 or less | 46 | 2 | 4.3 | <0.001 |
| 31-50 | 20 | 3 | 15.0 |  |
| >50 | 20 | 9 | 45.0 |  |
| Gender |  |  |  |  |
| Male | 32 | 4 | 12.5 | 0.46 |
| Female | 54 | 10 | 18.5 |  |
| Residence place |  |  |  |  |
| Durango State | 84 | 13 | 15.5 | 0.30 |
| Other Mexican State | 2 | 1 | 50.0 |  |
| Residence area |  |  |  |  |
| Urban | 79 | 12 | 15.2 | 0.72 |
| Suburban | 4 | 1 | 25.0 |  |
| Rural | 2 | 0 | 0.0 |  |
| Socio-economic level |  |  |  |  |
| Low | 65 | 9 | 13.8 | 0.25 |
| Medium | 15 | 4 | 26.7 |  |
| Educational level |  |  |  |  |
| No education | 26 | 6 | 23.1 | 0.35 |
| 1 to 6 years | 48 | 8 | 16.7 |  |
| 7-12 years | 11 | 0 | 0.0 |  |
| >12 years | 1 | 0 | 0.0 |  |

**Table 3.** Bivariate analysis of work factors and seroprevalence of HEV infection in waste pickers.

| Characteristic | Subjects<br>tested<br>No. | Prevalence<br>of<br>HEV<br>infection |  | *P*<br>value |
| --- | --- | --- | --- | --- |
|  |  | No. | % |  |
| Seniority |  |  |  |  |
| Up to 10 years | 44 | 3 | 6.8 | 0.03 |
| >10 years | 41 | 10 | 24.4 |  |
| Wearing gloves |  |  |  |  |
| Yes | 16 | 4 | 25.0 | 0.45 |
| No | 69 | 10 | 14.5 |  |
| Wearing face mask |  |  |  |  |
| Yes | 1 | 0 | 0 | 1.00 |
| No | 84 | 14 | 16.7 |  |
| Injuries at work |  |  |  |  |
| Yes | 61 | 10 | 16.4 | 1.00 |
| No | 23 | 4 | 17.4 |  |
| Eating during work |  |  |  |  |
| Yes | 48 | 10 | 20.8 | 0.21 |
| No | 37 | 4 | 10.8 |  |
| Eating from the garbage |  |  |  |  |
| Yes | 61 | 9 | 14.8 | 0.52 |
| No | 24 | 5 | 20.8 |  |
| Washing hands before eating |  |  |  |  |
| Yes | 64 | 11 | 17.2 | 1.00 |
| No | 21 | 3 | 14.3 |  |
| Alcohol consumption |  |  |  |  |
| Yes | 29 | 6 | 20.7 | 0.54 |
| No | 56 | 8 | 14.3 |  |

<sup>a</sup>Participants with available data.

**Table 4.**
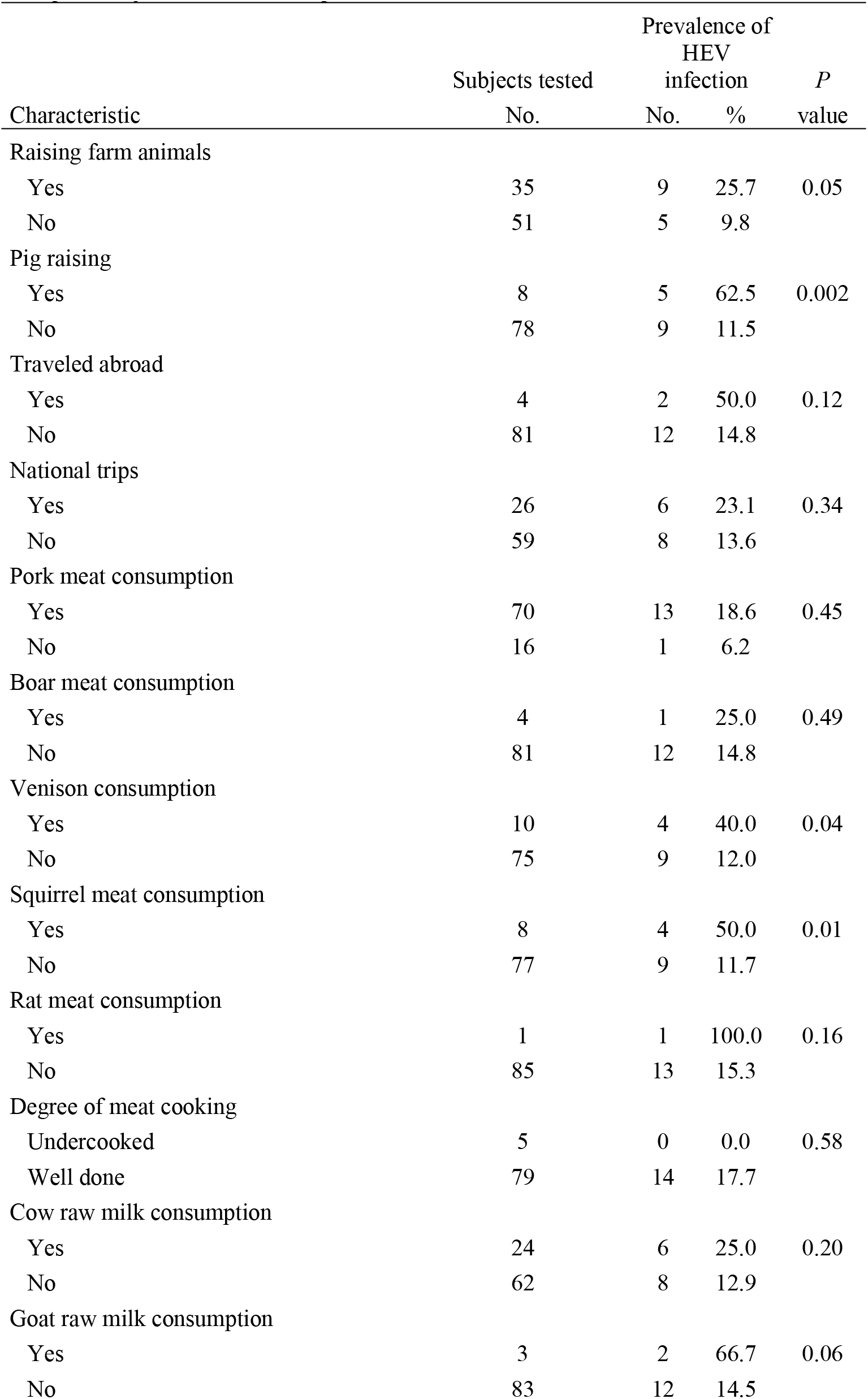
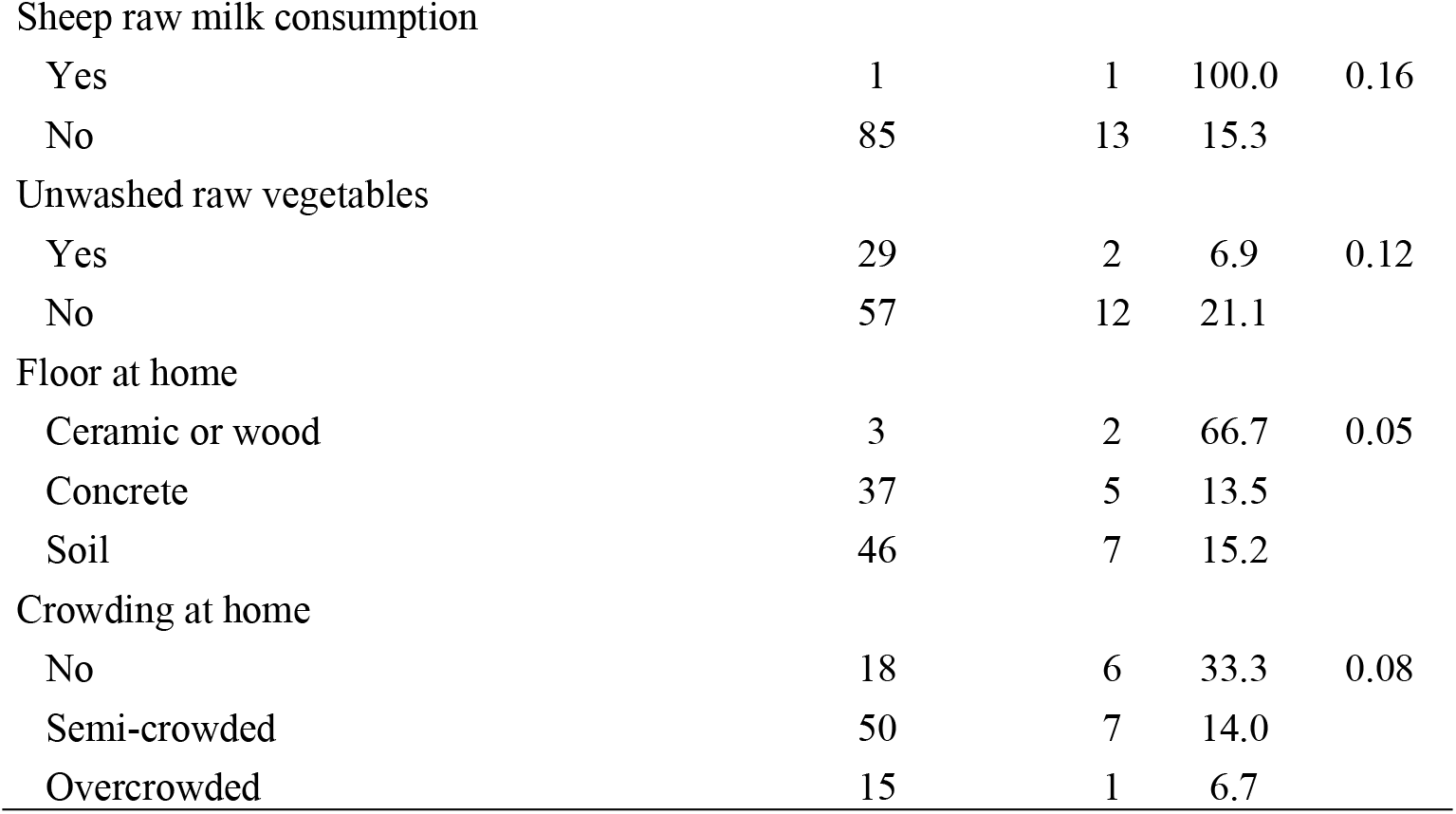
Bivariate analysis of a selection of behavioral and housing characteristics and seropositivity to HEV in waste pickers.

**Table 5.**
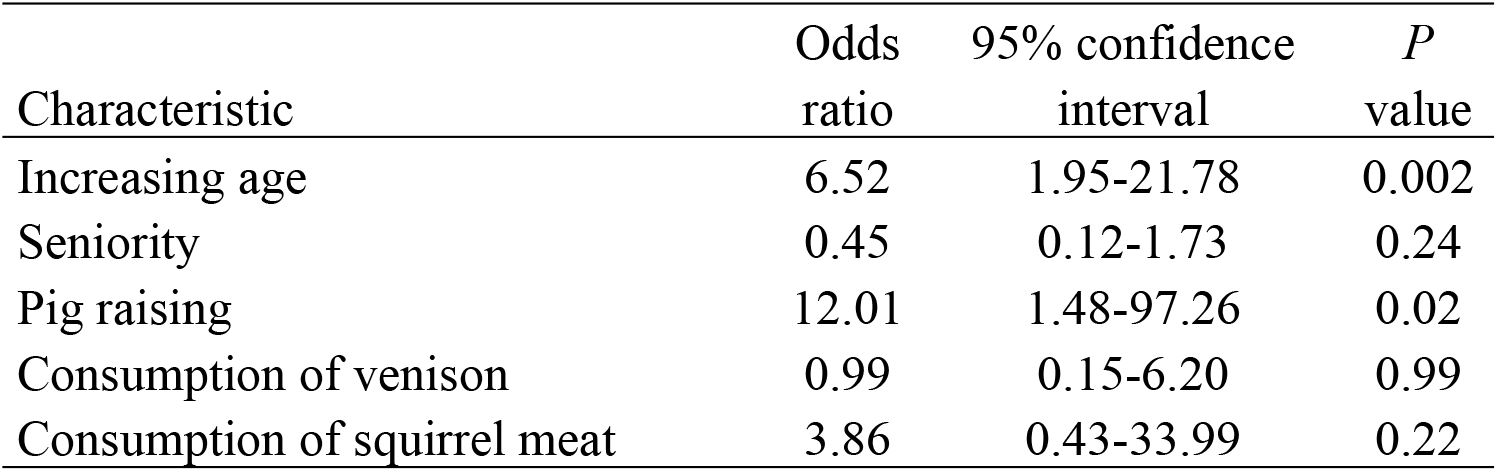
Multivariate analysis of selected characteristics of waste pickers and their association with HEV infection.

| Characteristic | Odds ratio | 95% confidence interval | *P* value |
| --- | --- | --- | --- |
| Increasing age | 6.52 | 1.95-21.78 | 0.002 |
| Seniority | 0.45 | 0.12-1.73 | 0.24 |
| Pig raising | 12.01 | 1.48-97.26 | 0.02 |
| Consumption of venison | 0.99 | 0.15-6.20 | 0.99 |
| Consumption of squirrel meat | 3.86 | 0.43-33.99 | 0.22 |

## Discussion

The seroepidemiology of HEV infection in waste wickers has been poorly studied. There has been only one previous study on the seroprevalence of HEV in waste pickers. In such study researchers reported the seroprevalence of HEV infection in waste pickers in a central Brazilian city [13]. The design of their study was cross-sectional, and the association between waste picker occupation and HEV seroprevalence was not determined. Therefore, in the present study, using an age and gender-matched case-control design, we aimed to determine the association between HEV seropositivity and the occupation of waste picker. Using the McNemar’s pair test, we found that waste pickers had a significantly higher HEV seroprevalence than controls. Thus, this result suggests that waste picker occupation is associated with HEV infection. The HEV seroprevalence of waste pickers was also associated with seniority in the activity by bivariate analysis. This result further supports the association between HEV seropositivity and waste picker occupation. The fecal oral route of HEV infection has been described as predominant in developing countries [3]. Garbage handled by waste pickers might be contaminated with HEV since fecal material is present in the city garbage. We further searched for sociodemographic, work, housing, and behavioral factors associated with HEV seroprevalence. Logistic regression analysis showed that only the variables increasing age and pig raising were associated with HEV seropositivity. Concerning the variable increasing age, our result agrees with previous observations that this factor is associated with HEV seropositivity in population groups in Mexico [15–17], and other countries [18–20]. With respect to the variable pig raising, our result is in line with previous observation about the link between HEV infection in humans and contact with pigs. In a meta-analysis of 32 studies from 16 countries, researchers found an increased seroprevalence of HEV infection in swine workers [21]. In another meta-analysis in China, researchers concluded that contact with pork or other pig products may be an important mode of HEV transmission [22]. In addition, a high seroprevalence of HEV infection in pigs in Durango, Mexico has been reported [23].

Intriguingly, ill waste pickers had a higher HEV seroprevalence than apparently healthy waste pickers. Similarly, waste pickers with reflexes impairment had a higher HEV seroprevalence than those without this impairment. We are not aware of a report about the link between HEV seropositivity and reflexes impairment. Acute and chronic HEV infections may lead to extrahepatic manifestations and has been associated with neurological diseases [24, 25], including for instance, neuralgic amyotrophy [26], acute encephalitis Parkinsonism [25], and Guillain Barré syndrome [27].

The limitations of the study include that no further testing for HEV infection as detection of IgM antibodies by enzyme immunoassay or DNA by polymerase chain reaction was performed; and participants were enrolled only in Durango City. Further research to determine the seroepidemiology of HEV infection in waste pickers should be conducted.

## Conclusions

This is the first age- and gender matched case-control study on the association between HEV infection and the occupation of waste picker. Waste pickers represent a risk group for HEV infection. Factors associated with HEV seropositivity found in this study may help in the design of optimal planning to avoid HEV infection.

